# Magellanic penguins balance navigation with foraging opportunities in complex current regimes

**DOI:** 10.1101/2024.12.12.628121

**Authors:** Richard M. Gunner, Flavio Quintana, Mariano H. Tonini, Mark D. Holton, Ken Yoda, Margaret C. Crofoot, Rory P. Wilson

## Abstract

Animals navigating in fluid environments often face lateral forces from wind or water currents that challenge travel efficiency and route accuracy. We investigated how 27 Magellanic penguins (*Spheniscus magellanicus*) adapt their navigation strategies to return to their colony amid regional tidal ocean currents. Using GPS-enhanced dead-reckoning loggers and high-resolution ocean current data, we reconstructed penguin travel vectors during foraging trips to assess their responses to variable currents during their colony-bound movements. By integrating estimates of energy costs and prey pursuits, we found that birds balanced direct navigation with current-driven drift: in calm currents, they maintained precise line-of-sight headings to their colony. In stronger currents, they aligned their return with lateral flows, which increased travel distance, but at minimal energy costs, and provided them with increased foraging opportunities. Since the lateral tidal currents always reversed direction over the course of return paths, the penguins’ return paths were consistently S-shaped but still resulted in the birds returning efficiently to their colonies. These findings suggest that Magellanic penguins can sense current drift and use it to optimize energy expenditure by maintaining overall directional accuracy while capitalizing on foraging opportunities.

## Introduction

Animals moving within water or air are subject to external fluid forces in the form of currents that can affect their locomotion, energy expenditure, and ability to navigate [see 1 for review]. These forces lead animals to drift away from heading-based trajectories, which poses navigational challenges for any extensive movements, such as during migration and foraging [1–5]. There is thus strong selective pressure for animals to detect and respond effectively to these environmental vectors [6, 7]. Several species have evolved strategies to mitigate the impact of currents by e.g. timing movements to coincide with favorable conditions [8, 9], adjusting speed [10, 11], or altering headings to compensate for drift [12–15]. However, the extent to which marine animals employ corrective mechanisms, which often navigate without visual landmarks or seabed references, is poorly understood [16]. A standard approach to assessing the effect of drift on animals is to study movements during goal-orientated traveling because the detrimental effects of fluid flow vectors can be examined directly. Such work shows, for example, that sea turtles have appreciable track tortuosity during migrations to their breeding islands [5], and similar deviations from supposed optimal trajectories have been observed in birds [17–19]. Explaining why these deviations occur and how animals maintain roughly goal-oriented behavior under environmental drift is hindered because we generally lack detailed information on the animal heading which, together with the external fluid vectors and travel speed, results in the observed path.

We used GPS-enabled dead-reckoning on breeding Magellanic Penguins *Spheniscus magellanicus,* central-place foragers operating out of sight of land during the breeding season [20–23], to study their abilities to return to their nests after foraging while exposed to considerable tide-driven current drift. Dead-reckoning combines information on the penguins’ compass-derived headings and modeled swimming speeds to reconstruct detailed movement paths [24]. This approach provided explicit information on how penguins adjusted their navigation strategies in response to variable ocean current conditions. By integrating estimated travel vectors (pre-current integration) with a hydrodynamic model of ocean currents, we quantified the complexity of their navigational challenges and reconstructed their responses to both minimal currents (slack water) and strong opposing flows. Additionally, we estimated energy expenditure using established power-speed relationships and analyzed dive profiles to assess prey pursuit behavior during their return journeys. These methods allowed us to explore how penguins balance efficient navigation with opportunistic foraging in a dynamic marine environment.

Our objectives were to: (1) investigate whether the penguins showed evidence of sensing current drift in the absence of landmarks (2) assess the penguins’ ability to orient toward their colony under minimal current conditions; (3) examine how they changed movement strategy when exposed to substantial currents; and (4) evaluate the energetic costs and distances traveled as a consequence of their navigation strategies.

## Materials and Methods

### Study site, subjects, instrumentation and attachment

Fieldwork was conducted between 22 November and 1 December 2019 at the San Lorenzo Magellanic penguin colony, Peninsula Valdés, Argentina (42.08° S, 63.86° W). We selected 27 adult Magellanic penguins (*Spheniscus magellanicus*) brooding small chicks for this study. Each penguin was equipped with a GPS logger (AxyTrek, Technosmart, Italy) and a Daily Diary (DD) logger [25]. The GPS units recorded positions at 1 Hz, while the DD loggers recorded tri-axial acceleration at 40 Hz and tri-axial magnetometry at 13 Hz, and pressure (=depth) at 4 Hz. Both devices were housed in hydrodynamic casings designed to minimize drag [26]. Combined, the devices weighed <1% of body mass and occupied <1% of cross-sectional area. Devices were secured on the dorsal midline with Tesa® tape [27] in a process taking less than 5 minutes. Penguins completed a single foraging trip before recapture.

Ethical approval for the research was granted by Swansea University’s Ethics Committee (SU-Ethics-Student: 260919/1894) and the Animal Welfare and Ethical Review Body (AWERB approval: IP-1819-30). Fieldwork permits were authorized by the Conservation Agency of Chubut Province (Disp N° 047/19-SsCyAP; No. 060/19- DFyFS-MP). All procedures involving penguin handling were reviewed and approved by the Dirección de Fauna y Flora Silvestre and the Ministerio de Turismo y Áreas Protegidas de la Provincia de Chubut.

### Ocean current model and dead-reckoning

Ocean currents were simulated using the Regional Ocean Modelling System (ROMS) [28], incorporating regional bathymetry and tidal constituents from the TPXO6 global model [29]. Model outputs were harmonically analysed for primary tidal components (M₂, S₂, and N₂), with U and V flow components representing east-west and north-south directions, respectively [30] (Text S1, Fig S1). The model output covered 50,500 km² at 1 km² resolution, spanning 22 November to 2 December 2019.

Penguin horizontal swimming speed was estimated based on their rate of change of depth and body pitch [31] using speed thresholds for surface swimming, low pitch underwater swimming, and high pitch diving. Penguins were dead-reckoned (DR) at 1 Hz resolution using the *Gundogs.Tracks()* function in R [32], corrected using GPS fixes acquired when the birds surfaced. At 1-second intervals, the U and V current components were bilinearly interpolated at nearest hourly timestamps according to the birds’ DR proximity using the *interpp()* function from the *akima* package in R. Outbound and inbound phases of each foraging trip were identified by cumulative changes in the shortest distance between the penguin and the colony, with the start of a continuous downward gradient marking the homing trajectory (Fig S3). Additional details on the ocean current model, horizontal swimming speed estimation, and adjustments for pressure sensor drift and tag orientation discrepancies are provided in Text S1.

### Heading and derivation of traveling vectors

Five, two-dimensional vectors were calculated based on bird or ocean current heading and speed (Fig 1):

**(i) *The ocean current vector:*** This represents the speed and direction of water movement from the ocean current model, with U (east-west) and V (north-south).
**(ii) *The real penguin travel vector relative to the water (pre-ocean current integration):*** This is the penguin’s travel vector as if it were unaffected by currents. It is calculated from the penguin’s heading (from the tag compass [30]) and speed, converted to U and V components. This vector represents the penguin’s movement relative to the surrounding water, prior to GPS dead-reckoning (DR) correction.
**(iii) *The real penguin travel vector relative to the ground (post-ocean current integration)*:** This describes the penguin’s effective movement across the ground, accounting for ocean current influence. It is computed by summing the bird’s vector relative to the water (see ii) and the ocean current vector (see i), providing U and V components of the penguin’s overall ground movement.
**(iv) *The optimal travel vector relative to the water*:** At each point along the DR track, we calculated the heading which, combined with the penguin’s speed (pre-DR correction and without ocean current integration), would produce a vector as close as possible to a direct ‘line-of-sight’ path back to the colony. If the penguin’s speed was insufficient to counter strong opposing currents, the ‘next best’ heading was selected to yield a travel vector nearest to the desired line-of-sight direction.
**(v) *The optimal travel vector relative to the ground*:** Calculated as the sum of U and V components from (i) and (iv). In most cases, this vector closely matched the line-of-sight heading.

Conversion between U and V components and speed and direction is detailed in Text S2.

**Figure 1.**
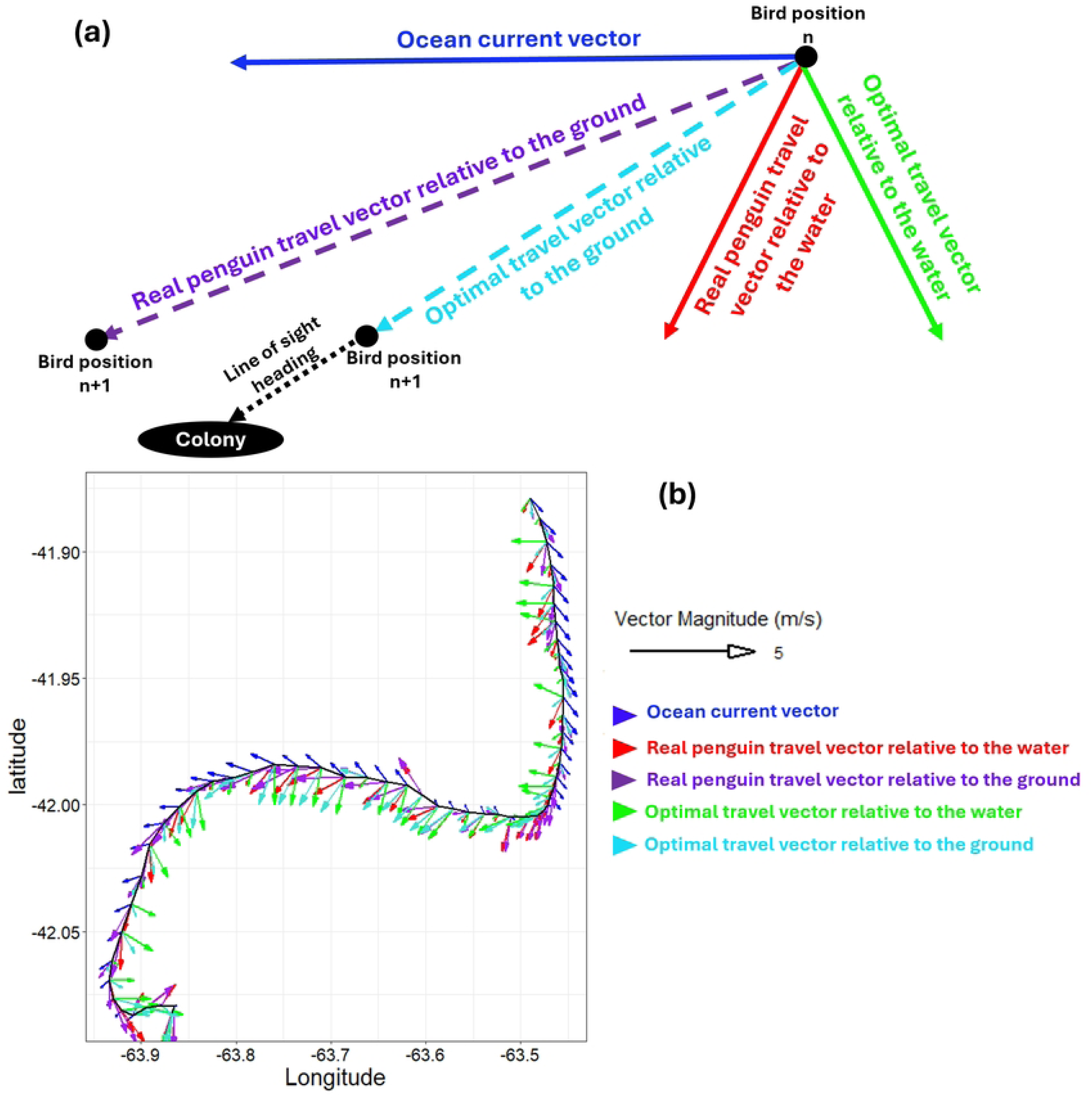
Key Vectors in Penguin Return Journey Analysis. This diagram illustrates the primary vectors used in analysing the penguin’s journey back to its colony. In panel (a), the ocean current vector (solid blue) combines with either the real penguin travel vector relative to the water (solid red) or the optimal travel vector relative to the water (solid green) at position ‘n.’ These interactions result in the projected real penguin travel vector relative to the ground (dashed purple) or the projected optimal travel vector relative to the ground (dashed cyan) at position ‘n+1’. The dotted black arrow represents the direct line-of-sight path to the colony. The optimal travel vector relative to the ground is aligned with this line-of-sight path unless the penguin’s speed is insufficient to fully counteract the current. Panel (b) shows a sample of these vectors recalculated every 5 minutes along a penguin’s southward return journey, based on its dead-reckoned track. The optimal travel vector relative to the water (solid green) sometimes counteracts the ocean current at angles greater than 90° to help the penguin stay on the most efficient path back to the colony.

### Ease of transport

The inverse of the cost of transport (COT) [cf. 33], defined as ‘ease of transport’ (the number of metres travelled per joule of energy (m/J), analogous to the ‘miles per gallon’ or ‘km per litter’ used by the vehicle industry), was used to emphasize how much the pathways utilized by penguins facilitated their progression within their moving oceanic landscape. Power requirements for the penguins’ chosen speed were calculated using the formula from Luna-Jorquera and Culik [34], relating mass-specific power to swim speed for the morphologically and physiologically similar congeneric Humboldt penguins (*Spheniscus humboldti*) [33].

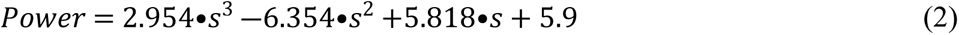

where *s* is the swimming speed (m/s) and *Power* is in W/kg. These values were then multiplied by 4 kg (the average weight of Magellanic penguins [23]) to convert to energy usage per penguin per second (J/s).

At each point along the DR track during the penguins’ inbound paths, coordinates were advanced based on the speed and direction of either the real penguin travel vector relative to the ground or the optimal travel vector relative to the ground. For each vector, two variants of ease of transport were calculated:

***(i)* The ‘ease of transport’ in any direction that reduces the distance to the colony:** This metric represents the energy efficiency of movement in any forward direction per unit power cost. It is calculated as the distance moved per second divided by the power cost (i.e., the distance between bird positions n and n+1 in Fig 1). Only movements within a 180° arc facing the colony were included; movements away from the colony were assigned an ease of transport value of zero.
***(ii)* The ‘ease of transport’ along the line-of-sight to the colony:** This metric represents energy efficiency of movement directly towards the colony, calculated as the distance covered along the line-of-sight trajectory to the colony per second, divided by power cost (i.e., the change in the line-of-sight distance between bird positions n and n+1 in Fig 1). Negative ease of transport values, arising when birds moved away from the colony, were set to zero.

### Agent-based model

We simulated penguin return journeys under a theoretical scenario where each penguin (agent) maintained a fixed initial line-of-sight heading toward the colony, without readjusting for cumulative current drift. Each agent used the real penguin’s speed data, iteratively applying its initial heading with time- and space-matched current vectors until either reaching the colony or failing to return to the colony’s longitude before the real penguin.

### Dive depth and prey pursuits

We determined the maximum dive depth of all dives (>0.3 m) in birds during their return journeys as well as an index of prey pursuit during dives based on ‘wiggles’ (undulations) in the depth profile following Simeone and Wilson [35]. Here, a single wiggle during which the penguin vertical velocity exceeded 0.3 m/s over ≥1 s was considered to represent a prey capture [35].

### Statistical analysis

#### Data binning and normalization

To address unequal sample sizes between penguins, we binned variables of interest, with counts normalized to proportions per penguin and averaged to yield grand mean relative frequencies. For analyses by distance travelled during return journeys, means were calculated per bin for each penguin and then averaged (Text S3).

#### Bootstrapped Kolmogorov-Smirnov (KS) test

A bootstrapped KS test was used to compare penguin heading deviations between outbound and inbound journeys under varying current strengths (’Slack’ < 0.3 m/s vs. ‘Appreciable’ ≥ 0.3 m/s), accounting for sample variability and intra-individual effects (Text S3). Approximately 30% of current data fell under the ‘Slack’ threshold, which was defined as less than 0.3 m/s—about half the mean current strength across all tracks. This threshold is below the penguins’ surface swimming speed, making it a logical boundary relative to their lowest observed travel speeds.

#### Generalized Additive Models (GAMs)

We employed GAMs (*mgcv* package in R) to investigate non-linear relationships between behavioural and environmental factors (see Text S3 for full details):

1. **Ease of Transport**: GAMs modelled the ease of transport relative to distance travelled, comparing real and optimal travel vectors, with random smooth effects to account for individual variation across the return distance (Text S3).
2. **Heading Deviation**: This GAM assessed factors influencing the deviation of the real bird travel vector heading from the line-of-sight to the colony. It included random smooth effects for individual variation across the return distance, with separate smooth terms for current speed, angular deviation between penguin and ocean current headings, resultant travel speeds (post- vs. pre-current integration), maximum dive depth, and rate of prey pursuits (Text S3, Fig S4).

## Results

### Penguin Movement Patterns

The majority of penguins exhibited ‘looping pathways’ [cf. 36], heading approximately northward to forage and returning southward to their colony. Return tracks often displayed an S-shape (Fig 2a), being influenced by the prevailing ocean currents which predominantly ran East-West or West-East depending on the phase of the tidal cycle (Fig 2b). On average (mean ± SD), the return journeys began 50 ± 15 km from the colony and took 12 ± 4 hours to complete, whereas outbound journeys lasted 20 ± 9 hours. Penguins exposed to cross-currents during both phases spent approximately equal time in eastward and westward currents. During the outbound phase, they spent 50.7% ± 10.5% of the time in eastward currents and 49.3% ± 10.5% in westward currents [(paired t-test: t = −0.366, p = 0.717)]. Similarly, during the inbound phase, they spent 49.3% ± 14.6% of the time in eastward currents and 50.7% ± 14.6% in westward currents [(paired t-test: t = 0.237, p = 0.814)]. The average current strength across all tracks was similar between the two phases, averaging 0.59 ± 0.23 m/s during the outbound phase and 0.60 ± 0.20 m/s (max 2.0 m/s) during the inbound phase (cf. Fig S5). These represent a significant fraction (mean - 29%, max 95%) of normal penguin travelling speed of *ca*. 2.1 m/s [37, 38] and should have a corresponding influence on their trajectories. Birds landed on the coast at minimal distance from their departure point at the colony (0.17 ± 0.18 km, range: 0.01–0.72 km), with 85% of birds returning within 0.3 km.

**Figure 2.**
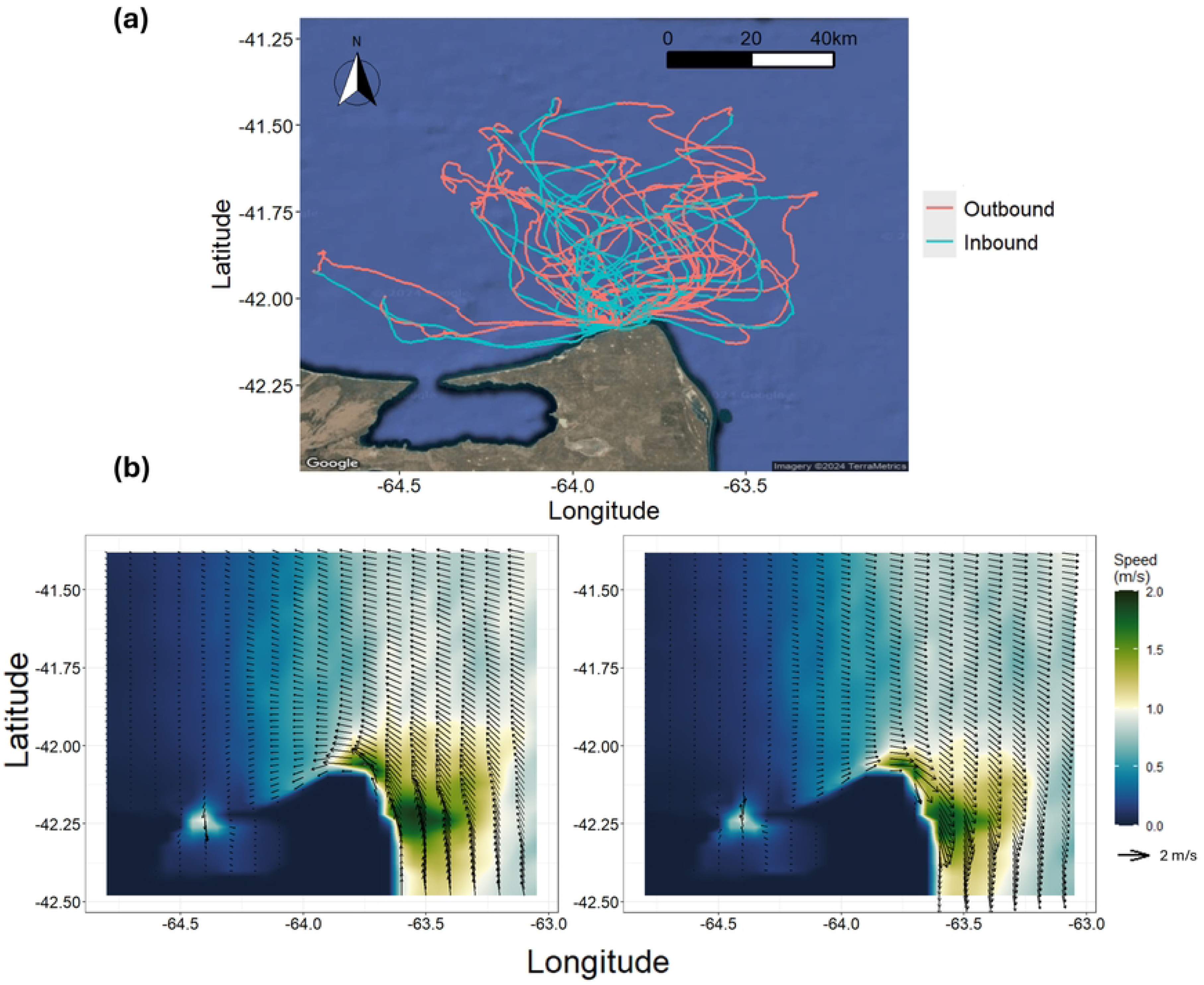
Tracks of 27 Penguins and Visualized Current Conditions. (a) The GPS- corrected dead-reckoned tracks of 27 penguins at sea, coloured to distinguish between the outbound (red) and inbound (blue) phases of their foraging trips. (b) The current conditions within the grid area, showing peak tidal strength during a single day as a function of the tidal cycle, including the outgoing (‘ebb’ - left panel) and incoming (‘flood’ - right panel) tides. Penguins are often subjected to strong cross-currents during these phases.

### Influence of currents on heading deviation

During inbound journeys, penguin travel vector headings relative to the water were more tightly distributed around their line-of-sight heading than during outbound phases (D = 0.232, 95% CI: 0.228–0.241; Fig 3). The variability of heading strategies was significantly influenced by current strength and direction (Table 1). During inbound movement, on average, penguins directed themselves within 25° of the line-of-sight heading 60% of the time and within 45° 80% of the time, rarely deviating more than 90° regardless of current conditions (Figs 3 and S2).

**Figure 3.**
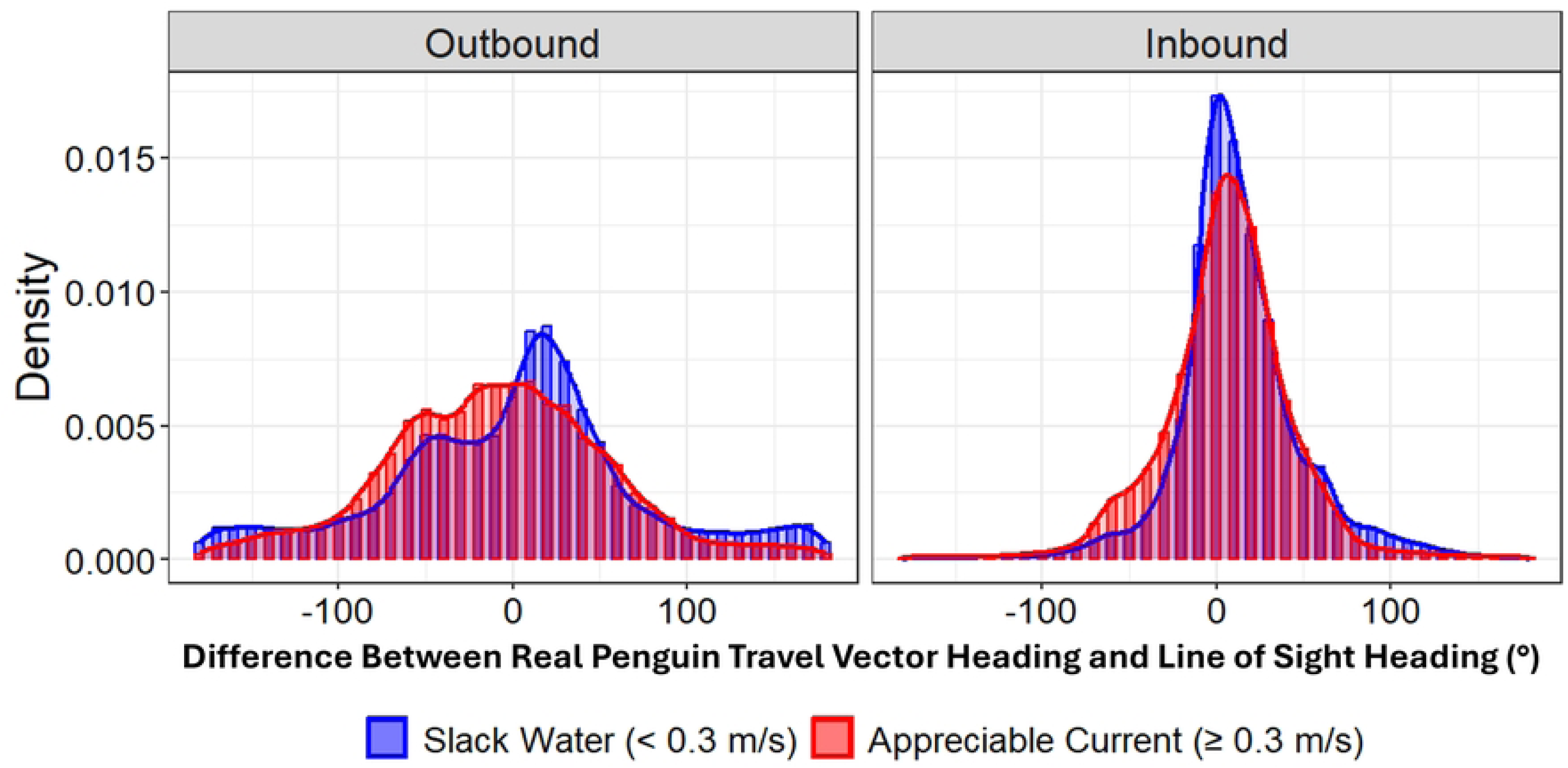
Penguin Heading Distributions During Outbound and Inbound Phases of Foraging Trips Under Different Current Conditions. This plot shows the difference in relative density distribution between the penguin’s travel vector heading (relative to water) and the line-of-sight heading. In both ‘Slack Water’ (< 0.3 m/s, blue) and ‘Appreciable Current’ (≥ 0.3 m/s, red) conditions, penguins align more closely with the line-of-sight heading during the return (inbound) phase, suggesting a more focused return to the colony compared to the departure (outbound) phase. A heading difference of 0° indicates direct movement toward (inbound) or away from (outbound) the colony, while 90° indicates perpendicular movement.

**Table 1.**
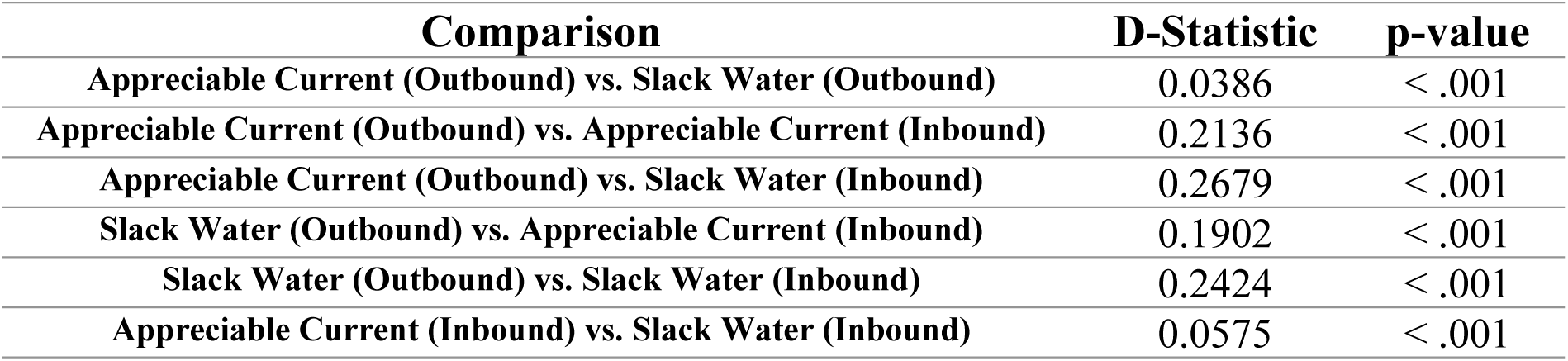
Pairwise Kolmogorov-Smirnov (KS) Test Results for Penguin Heading Distributions Relative to the Line-of-Sight heading. Comparisons are made for real penguin vector headings relative to the water across different ocean current conditions (’Slack Water’ < 0.3 m/s and ‘Appreciable Current’ ≥ 0.3 m/s) and trip phases (Outbound vs. Inbound). The D statistic represents the maximum difference between the cumulative distributions of the two compared groups, with corresponding p-values indicating significance.

To understand how ocean currents affect penguin navigation during the inbound phase of their foraging trips, we analysed the relationship between penguin travel vector headings and their line-of-sight direction to the colony in relation to ocean current direction.

Before accounting for ocean current effects, the real penguin travel vector headings relative to the water (represented by the red arrow in Fig 1) sometimes required the penguins to swim against the ocean current to maintain their line-of-sight heading (dotted black arrow in Fig 1) (see Fig 4a). When ocean current vectors were included, the resultant penguin travel vector headings relative to the ground (purple arrow in Fig 1) showed more pronounced deviations from this direct line-of-sight path (see Fig 4b). In contrast, theoretically optimal travel vector headings relative to the water, calculated before integrating ocean current effects (green arrow in Fig 1), would often require the penguins to swim at angles greater than 90° from the ocean current direction to achieve a resultant travel vector heading leading directly back to the colony (cyan arrow in Fig 1) (see Fig 4c and Fig 4d).

**Figure 4.**
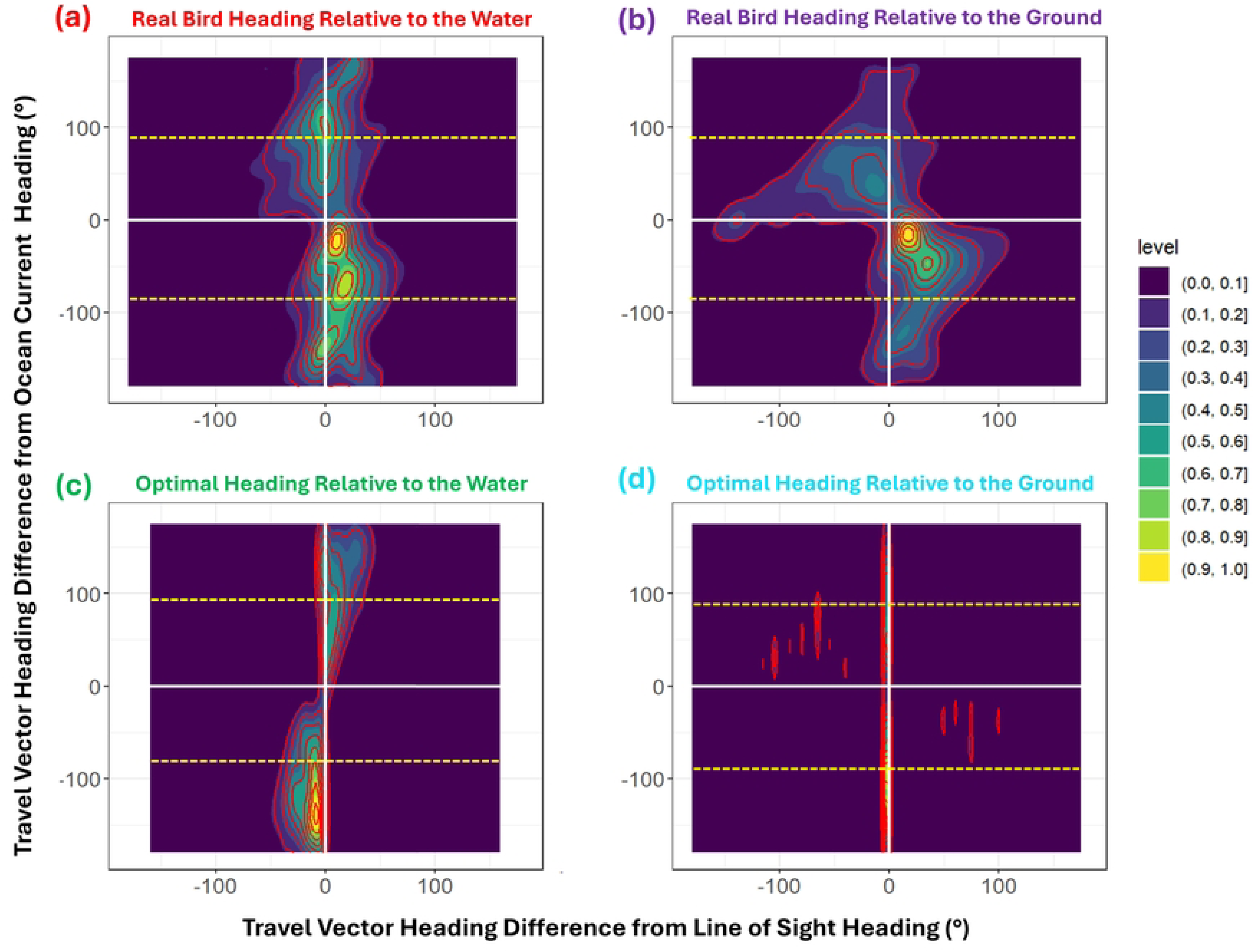
2D-Binned Contour Plots of Real vs. Theoretically Optimal Penguin Travel Vectors During the Return Phase of Foraging Trips. These contour plots compare real and theoretically optimal penguin travel vectors relative to the line-of-sight heading to the colony (X-axis) and the ocean current direction (Y-axis). *(a)* Real penguin travel vector headings relative to the water generally align with line-of-sight trajectories toward the colony, though some skew is evident. *(b)* When accounting for ocean current effects, the resultant real penguin headings relative to the ground reveal current-assisted trends, roughly indicated within the dashed yellow lines. These headings show increased dispersion around the line-of-sight heading, with more occurrences of nearly opposing headings. *(c)* The theoretically optimal penguin travel vector headings relative to the water, which ideally allow the birds to follow a direct line-of-sight path back to the colony after factoring in ocean current effects (as seen in panel d), would often require significant time swimming against the current, especially outside the dashed yellow lines.

Variation in penguin travel vector headings around the line-of-sight heading differed notably among individuals and changed non-linearly with distance to the colony (Fig 5a). Penguins showed a general trend toward more goal-oriented behavior at the start and end of their return journeys, with greater deviations in the middle. This pattern coincided with variations in the proportion of time spent swimming against ocean currents, especially under appreciable current strength (Fig 5b).

**Figure 5.**
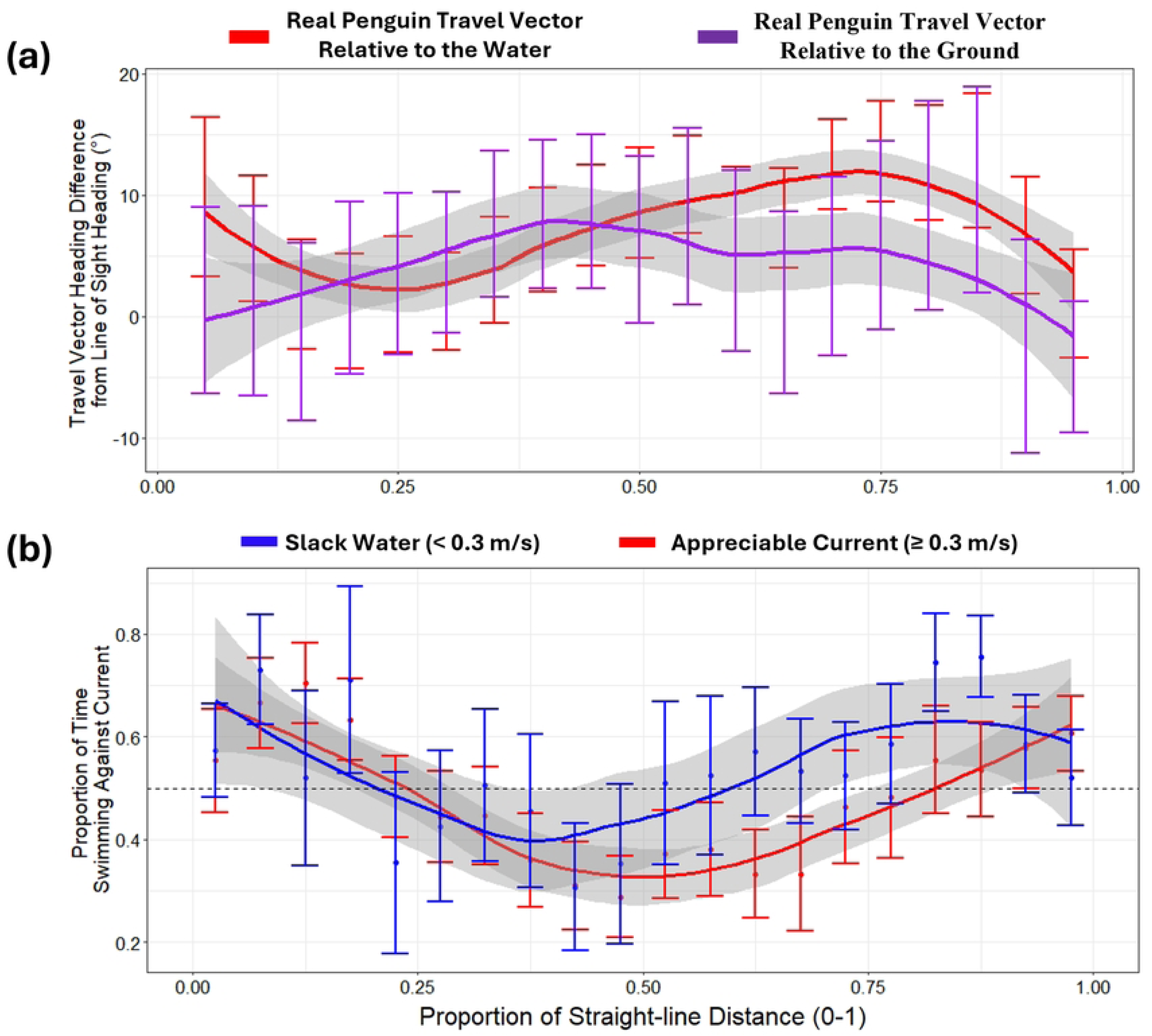
Variation in Penguin Headings Relative to Line-of-Sight and Proportion of Time Spent Swimming Against Ocean Currents During the Return Journey. (a) Mean (± 1 SE) angular difference between real penguin travel vector headings and the line-of-sight direction to the colony. Headings are shown relative to the water (before accounting for ocean currents; red) and relative to the ground (after accounting for ocean currents; purple), with fitted loess smooth lines. (b) Mean (± 1 SE) proportion of time penguins spent swimming against the ocean current. Blue represents periods with “slack water” (< 0.3 m/s), while red indicates periods with “appreciable current” (≥ 0.3 m/s). Time spent swimming against the current is defined as instances where heading angles exceeded 90° in absolute terms. The dashed horizontal line at y = 0.5 marks the 50% threshold, indicating equal time spent swimming with and against the current. In both panels, the x-axis is divided into 0.05 increments of straight-line distance travelled.

### Ease of Transport

Penguins achieved higher ease of transport values if they moved generally, but not directly, towards the colony by following their real travel vector headings relative to the ground, compared to the theoretical optimal travel vector scenario (Fig 6a). For movement along the line-of-sight path, ease of transport was generally slightly lower at the start of the return journey, aligning more closely with the optimal travel vectors in the middle section, a pattern that coincided with increased time spent swimming with ocean currents during this part of the journey (Fig 5b).

**(i) The ease of transport in in any direction that reduces the distance to the colony:** The GAM analysis indicated that the theoretical optimal travel vectors had a significantly lower ease of transport than the real penguin travel vectors across the return journey (Estimate = −0.0054, SE = 0.0005, p < 0.001; Fig 6a, Text S3, Table S1). Pairwise comparisons showed that this significance spanned the majority of the return journey (Fig 6a).
**(i) The ease of transport along the line-of-sight to the colony:** The GAM analysis indicated that the theoretical optimal travel vectors had a slightly higher ease of transport along the line-of-sight path than the real penguin travel vectors (Estimate = 0.0014, SE = 0.0005, p = 0.0068; Text S3, Table S1). Pairwise comparisons revealed that this significant difference was limited to the early phases of the return journey (Fig 6a).

**Figure 6.**
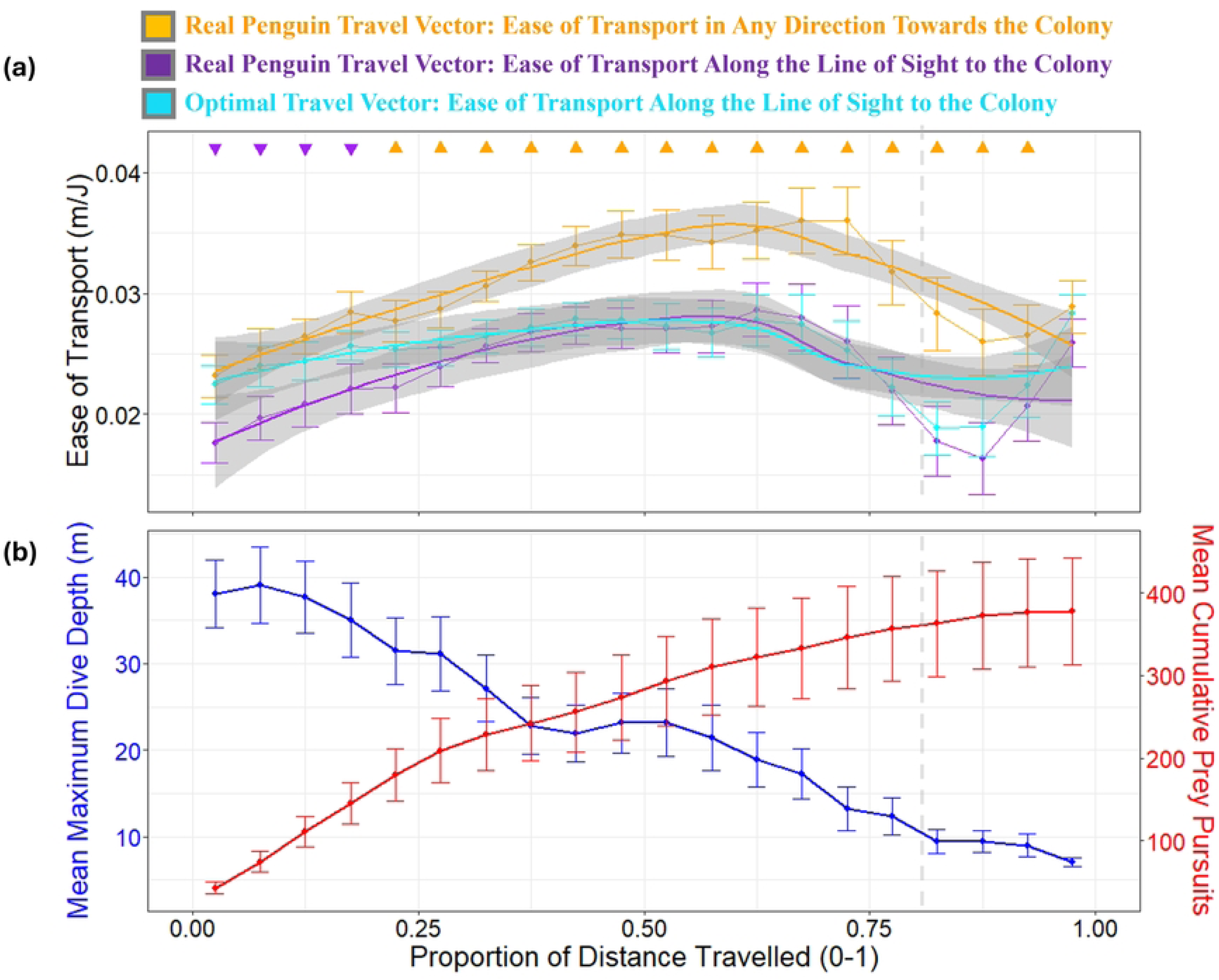
Penguin Ease of Transport for Real vs. Optimal Travel Vectors Relative to the Ground During the Return Journey with Respect to Depth use and Prey Acquisition. (a) Mean ease of transport (± 1 SE, in m/J) is shown across the proportion of the return distance to the colony for the real penguin travel vector in any direction towards the colony (orange), the real penguin travel vector along the line-of-sight to the colony (purple), and the optimal travel vector (cyan). Higher values indicate greater energy efficiency of movement. Significant pairwise differences (p < 0.05) between the real and optimal vectors are marked with solid triangles at 0.05 intervals of the return journey, based on GAM-predicted ease of transport values, which account for individual variability and smooth trends over distance (see Text S3 for details). Upward-facing triangles indicate that the real travel vector (orange or purple) had significantly higher ease of transport than the optimal vector (cyan), while downward-facing triangles indicate the opposite. The absence of a triangle indicates no significant difference at that interval. (b) Mean maximum dive depth (± 1 SE, in m) and the estimated mean cumulative number of prey items captured during the return journey, plotted at 0.05 intervals of the return journey’s proportion of distance travelled. Note that the overall depth decreases over time showing that birds allocate increasing time to horizontal travel. Dive depths decrease to less than 10 m when ca. 80% of the journey is complete (marked by the dashed grey vertical line). This pattern corresponds with substantive changes in the ease of transport and minimal additional prey capture. Importantly, this reduction in dive depth over time was not influenced by the surrounding seafloor depth (Fig. S6).

### Agent-based model

The ‘what if’ scenario where the line-of-sight heading was not corrected over time resulted in many simulated birds returning to land at great distances from their breeding colony (S7 Fig)

### Heading deviation from line-of-sight

The GAM revealed that deviation of bird vector headings relative to the water from the line-of-sight heading was significantly influenced by individual differences, trip progression, and environmental factors. Penguins adjusted their navigation strategies along the return journey, with significant effects observed for the angular difference from the ocean current, maximum dive depth, and resultant speed after current integration. Additionally, prey pursuits influenced heading deviation, and a complex interaction among ocean current speed, alignment with ocean current direction, and resultant bird vector speed differences also impacted heading deviation (Text S3, Fig S4).

### Foraging

Penguin continued to dive to depths in excess of 10 m and likely catch prey [35] up to ∼80% of their return journey (Fig 6b and Fig S8).

## Discussion

### Adaptation to current conditions and goal-oriented navigation

Magellanic penguins must navigate efficiently over long distances to return to their nests and feed their chicks. Our findings show that they often orient themselves toward the colony by following a general line-of-sight heading (Figs 4 and S9), especially under calm ocean current conditions (Fig 3). This suggests an innate, goal-oriented navigation ability that is effective even without visible landmarks [39–41]. Given the penguin’s low profile and minimal visibility at the sea surface, visual contact with the colony is impossible or unlikely for most of the journey, implying reliance on alternative sensory mechanisms. Penguins may use geomagnetic cues to detect the Earth’s magnetic field for orientation, as magnetoreception has been suggested in other avian species [42]. Mechanoreception—detecting water flow or pressure changes across their bodies, might also help gauge current direction and strength [cf. 43]. These mechanisms would enable them to maintain their course effectively even in dynamic marine environments.

Interestingly, penguins tend to navigate more directly at the beginning and end of their return journeys, with greater heading deviations occurring mid-journey, particularly in stronger currents (Fig 5). This pattern suggests an adaptive strategy to balance direct navigation with environmental drift (*cf.* Figs S10 and S11). Closer to the colony, penguins show a stronger tendency to follow the line-of-sight, which may indicate reliance on visual cues near the colony, similar to behaviours observed in shearwaters [44]. However, 8 of 27 penguins returned at night (10 p.m. to 4 a.m.), indicating that visual cues may not be essential for their homing [cf. 44]. Deviations from direct trajectories become increasingly detrimental as the colony approaches, reinforcing the importance of direct navigation in the final stages of the journey. Impressively, 85% of penguins returned within 0.3 km of their departure points, matching the accuracy reported by Quintana, Gómez-Laich [45]. To illustrate, for an average 50 km journey, a 300 m deviation represents a 99.4% direct return efficiency.

### Flow-assisted navigation and adaptive heading adjustment for energy efficiency

The tidal cycles in the San Matías Gulf expose penguins to alternating east-west currents in a semi-diurnal cycle (approx. 12.4 hours), causing lateral drift but also presenting opportunities for current-assisted movement (Figs 2, S1, S2, S10). Given the average 12-hour duration of their return journeys, penguins typically encounter both current directions in approximately equal measure (Fig S5), allowing the current regime to correct for itself over time without continuous effort by the penguins to work against the currents. Our observations on the precise headings taken by the penguins certainly suggests they can sense current vectors, as evidenced by their differences in headings during slack water and stronger currents (Fig 3), even when out of sight of land (e.g., Figs S10 and S11). In addition, it seems that they specifically allow lateral drift because it can later be corrected by reversed currents (e.g., Fig 1 and S10). Specifically, they adjust their headings to avoid opposing currents and take advantage of favourable flows, when possible, sometimes angling their path relative to the line-of-sight to maintain forward progress while conserving energy (e.g., Figs 5, S10, S11). This flow-assisted movement behaviour [1] is evident in the ease of transport values: penguins’ real travel vectors achieved higher efficiency relative to the ground than the theoretical optimal vectors (which prioritize the most direct movement back to the colony) with birds nonetheless decreasing the distance between themselves and the colony (orange line in Fig 6). By harnessing current strength (Figs 4, 5, S2, S9, S11), penguins enhanced their travel efficiency in a manner similar to birds using tailwinds during migration [7, 46].

Swimming with the current during mid-journey stages reduces drag-related energy costs and conserves energy compared to swimming directly against currents [47–49], which is crucial given the exponential increase in power costs at higher swimming speeds (see eqn 2). We suggest that this strategy also allows penguins to search for prey opportunistically because dive data reveal frequent deep dives during the return trip where penguins captured prey (Fig 6b, S4, S8, S11), even though surface swimming typically offers the most energy-efficient travel for air-breathing marine animals [50]. This adaptive strategy highlights the penguins’ flexibility in heading adjustment to balance energy conservation with steady progress toward the colony. Notably, the ease of transport along the line-of-sight to the colony was only slightly higher overall in the theoretical optimal scenario (cyan vs. purple lines in Fig 6a), with penguins’ real movements achieving comparable efficiency despite deviating from a direct line-of-sight heading (Figs 4b and S9). In other words, by adjusting their headings to take advantage of favourable currents, penguins can maintain efficient progress toward their goal, even if their path deviates from the most direct route.

### Navigation strategies and environmental complexity

The variable tidal flows in the San Matías Gulf present a dynamic and challenging environment for navigation. Given the eastward and westward currents they encounter, why don’t penguins simply maintain a general line-of-sight heading without compensating for drift? Our results indicate that penguins actively correct for current-induced drift in a non-linear manner, readjusting to a ‘line-of-sight’ heading when currents allow flow-assisted movement (i.e., Fig S7a *vs.* S7b). This navigation strategy is complex, as shown by our GAM analysis, which revealed that deviations from the line-of-sight heading are influenced by multiple factors, including current speed, alignment with the current, opportunities for prey pursuits, and dive behaviour (Text S3, Table S2, Fig S4). This variety of influences underscores the intricate nature of penguin navigation in an ever-changing foraging environment, and indicates that their movement strategies are complex, adapting over current regimes, time, space, and in response to prey availability. We suggest that similar complexities may occur in animal navigation challenges elsewhere and that these may account for departures from expected routes [5].

## Acknowledgments

We would like to express appreciation to the individuals from Ea. San Lorenzo for logistical support, to the Instituto de Biología de Organismos Marinos (IBIOMAR) – CONICET, and the CCT CENPAT-CONICET for institutional and logistical assistance, and to the Ministerio de Desarrollo Territorial y Sectores Productivos and the Secretaría de Turismo de la Provincia de Chubut, Argentina, for granting permits to work at the Península Valdés natural protected area.

## Supporting information

**Text S1.** Numerical Ocean Current Model, Dead-Reckoning Speed Estimates, and GPS-Corrected Dead-Reckoning Procedure.

**Text S2.** Conversions Between U and V Components (m/s) and Heading (°) and Speed (m/s).

**Text S3.** Statistical Methods Expanded

**Fig S1.**
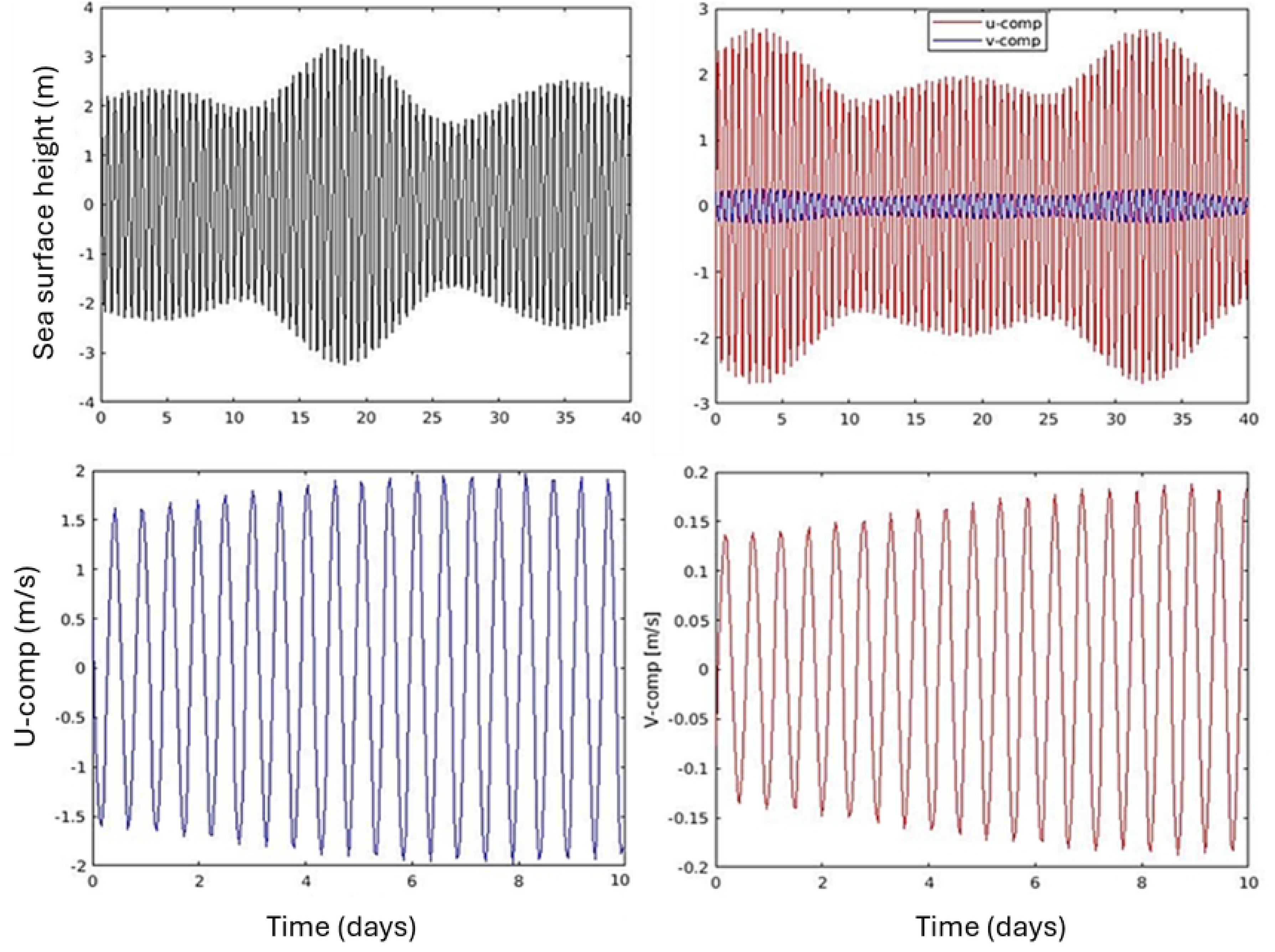
Time Series of Elevation and Ocean Current Components Highlighting Cross-Current Variations. The **Top Panel** shows the variation in elevation and the U (east-west) and V (north-south) components of ocean currents in the study region over a 40-day period. The **Bottom Panel** presents a zoomed-in view of a 10-day section focusing on the phases of the U and V current components. The plot demonstrates periods dominated by cross-currents, particularly highlighting the U component’s influence on ocean dynamics.

**Fig S2.**
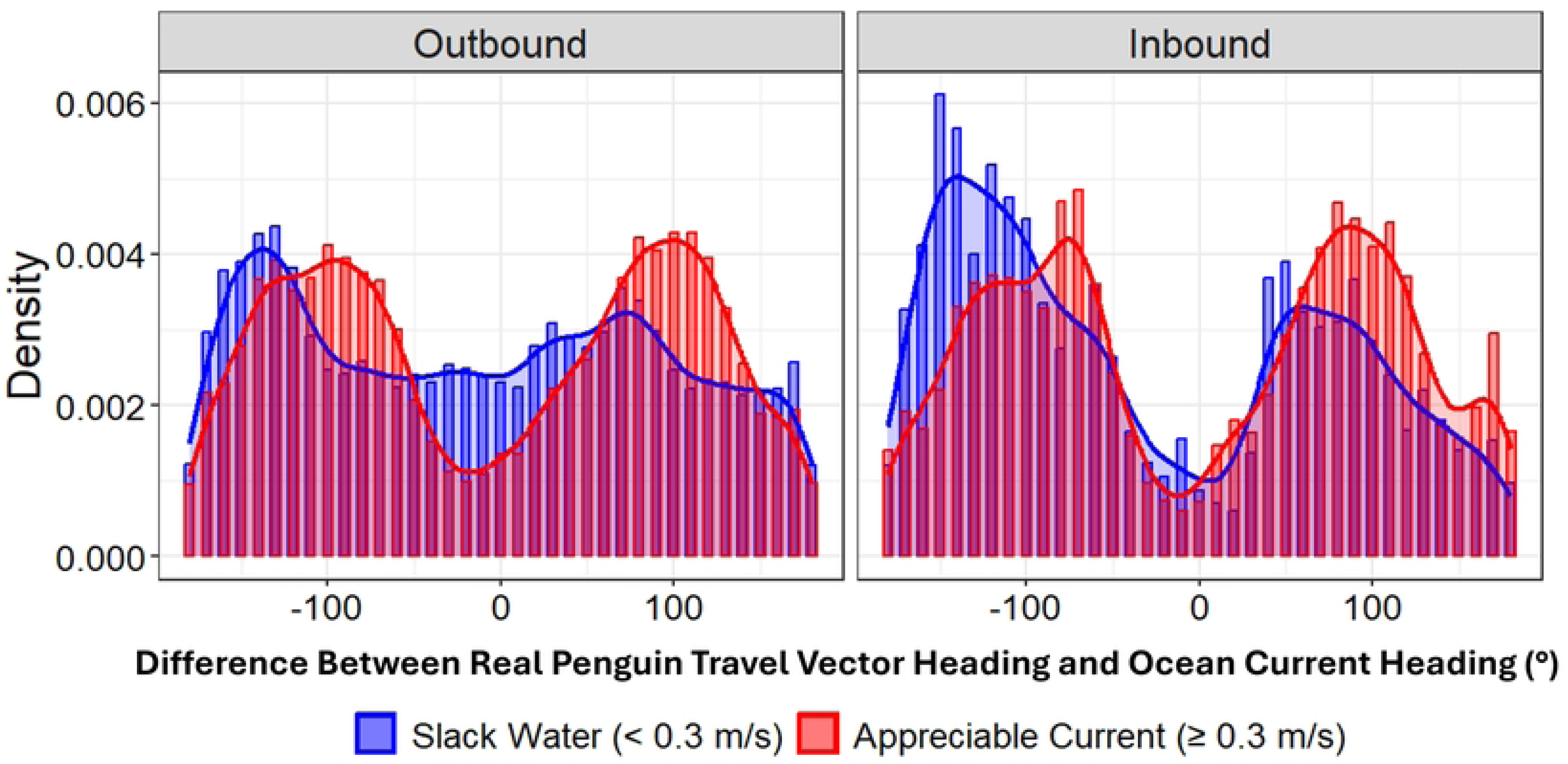
Penguin Heading Distributions During Outbound and Inbound Phases of Foraging Trips Under Different Current Conditions. This plot shows the difference between the penguin’s travel vector heading (relative to the water) and the ocean current direction. The distribution is often bimodal, likely due to encounters with cross-currents. This bimodality is more pronounced during the return phase, suggesting more frequent cross-current navigation.

**Fig S3.**
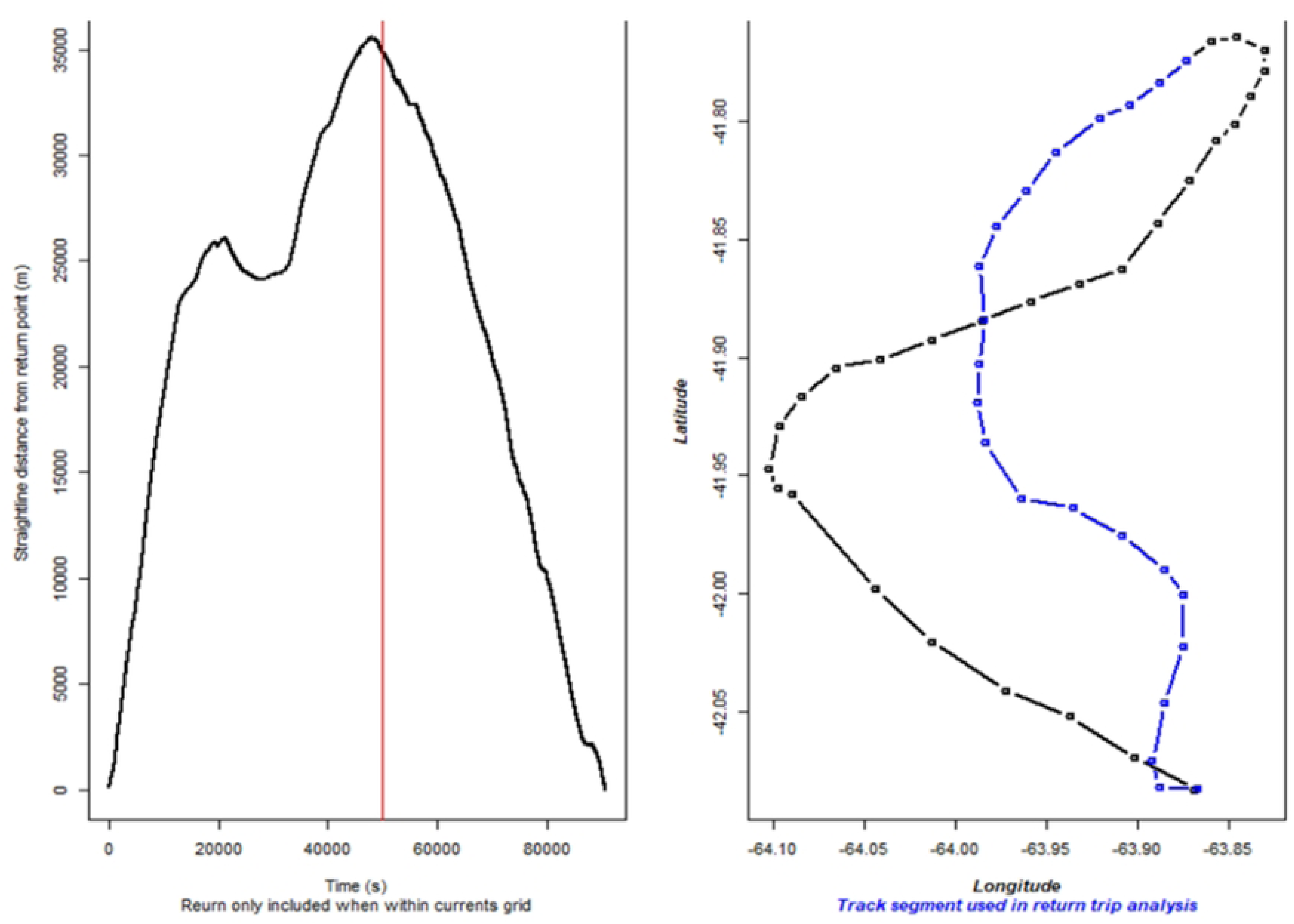
Determining the Start of a Penguin’s Return Journey Using GPS Data and Distance to Colony. The **Left Plot** depicts the shortest straight-line distance from the penguin to the colony over time. The red vertical line indicates the start of the return phase, determined by selecting all data after the first GPS fix that occurred during the beginning of a continuous downward trend in straight-line distance to the colony. The **Right Plot** shows the penguin’s movement track (15-minute GPS fix intervals shown). The blue section of the track represents the ‘Inbound’ phase, corresponding to the data after the red vertical line in the left plot.

**Table S1. Summary of GAM Results for Ease of Transport Models:**

**i. The real penguin travel vector relative to the ground.**

**ii. The optimal travel vector relative to the ground.**

**Table S2. Summary of GAM Results for Deviation of Penguin Heading from Line-of-Sight Heading.**

**Figure. S4.**
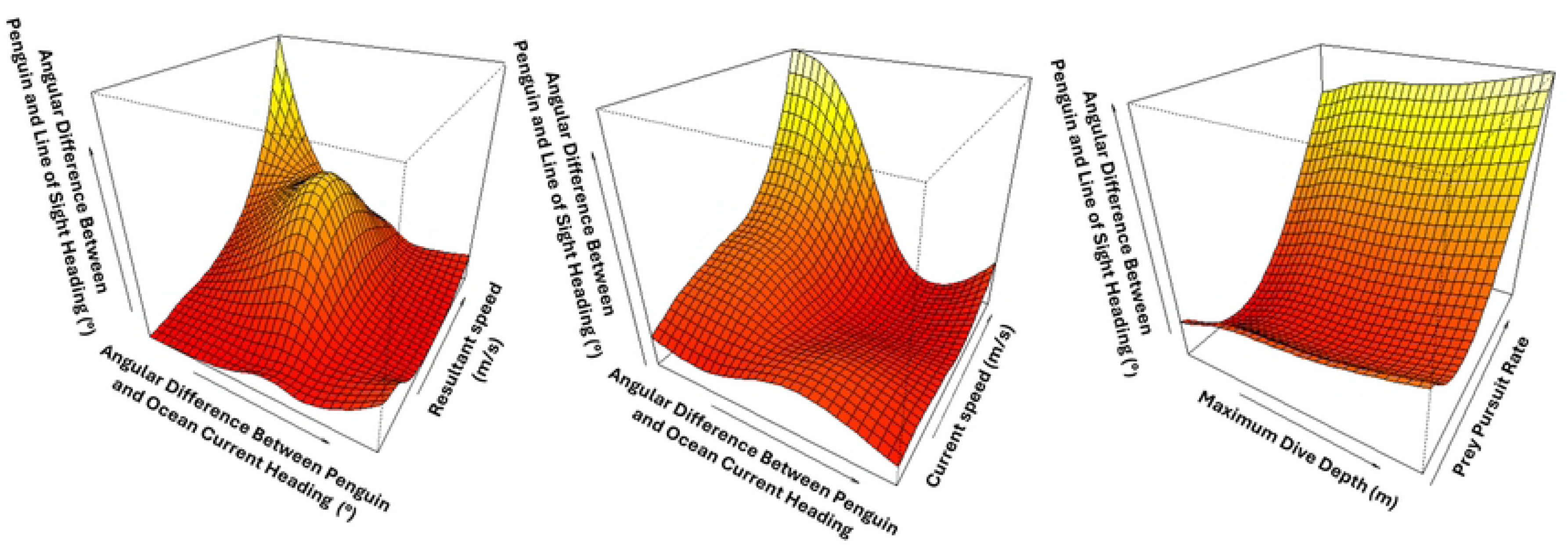
Three-dimensional plots generated using ‘vis.gam’ provided insights into how pairs of predictors interact to influence penguins’ deviation from their intended line-of-sight direction (cf. Table. S2). (a & b) Due to the nature of cross-currents in the region, penguins tend to align their movement with ocean current flow vectors. This alignment results in greater deviation from their line-of-sight trajectory, particularly at higher current speeds. However, this alignment also enables penguins to travel greater distances per unit time by increasing their resultant speeds. *(c)* Penguins’ heading deviation from the line-of-sight direction generally increases in a non-linear fashion with respect to increasing dive depth and prey pursuit rate.

**Figure S5.**
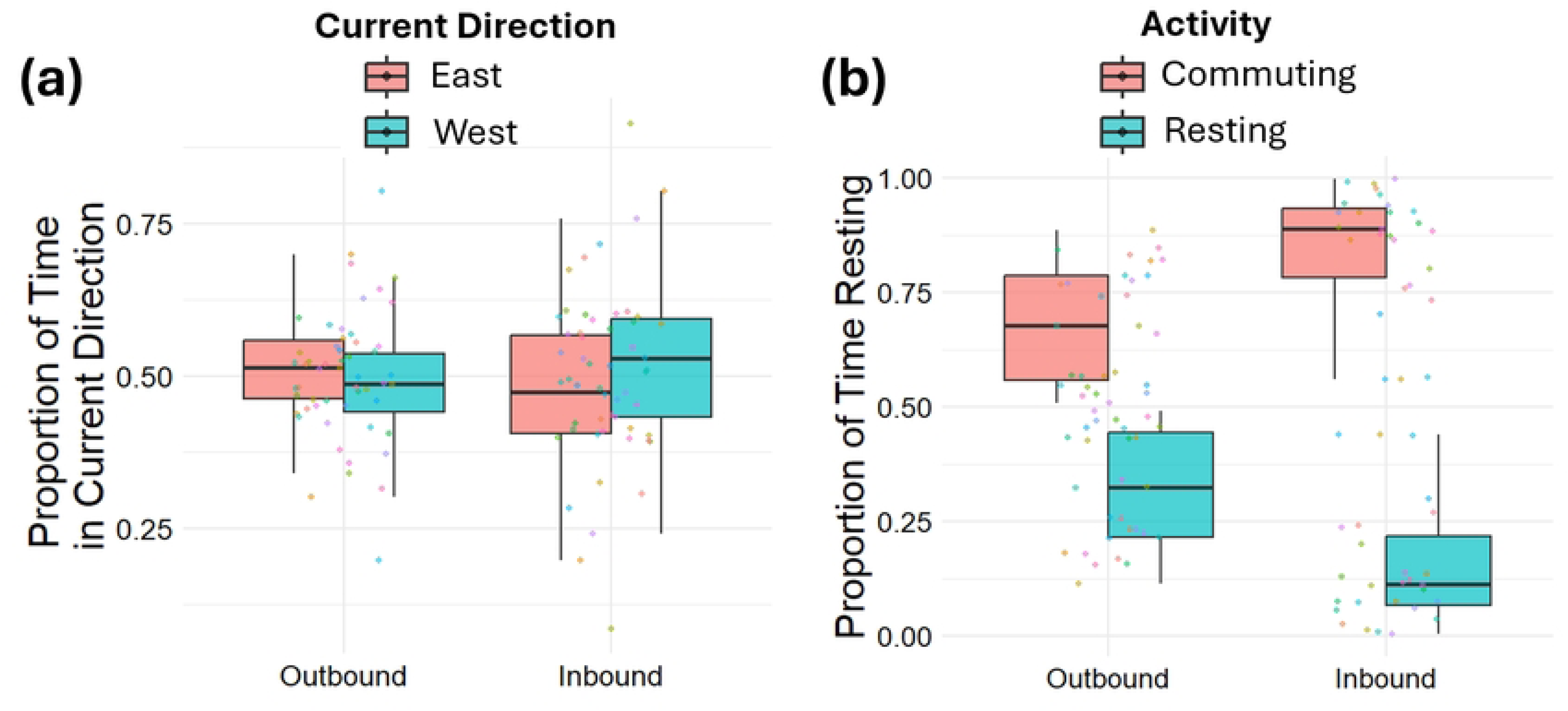
Proportion of Time Spent in Eastward vs. Westward Currents and Activity Types During Outbound and Inbound Trips. (a) Shows the proportion of time penguins experienced eastward-facing and westward-facing currents during outbound and inbound trips. (b) Displays the proportion of time spent in each activity type (resting vs. commuting) during outbound and inbound trips. Resting was defined as surface periods ≥ 60 s and commuting-specific to this figure-as all other times. Boxplot (boxes encompass the 25∼75 % interquartile range, horizontal bars reflect the median and whiskers extend to 1.5 * Interquartile range). Jittered points show each bird’s data point.

**Figure S6.**
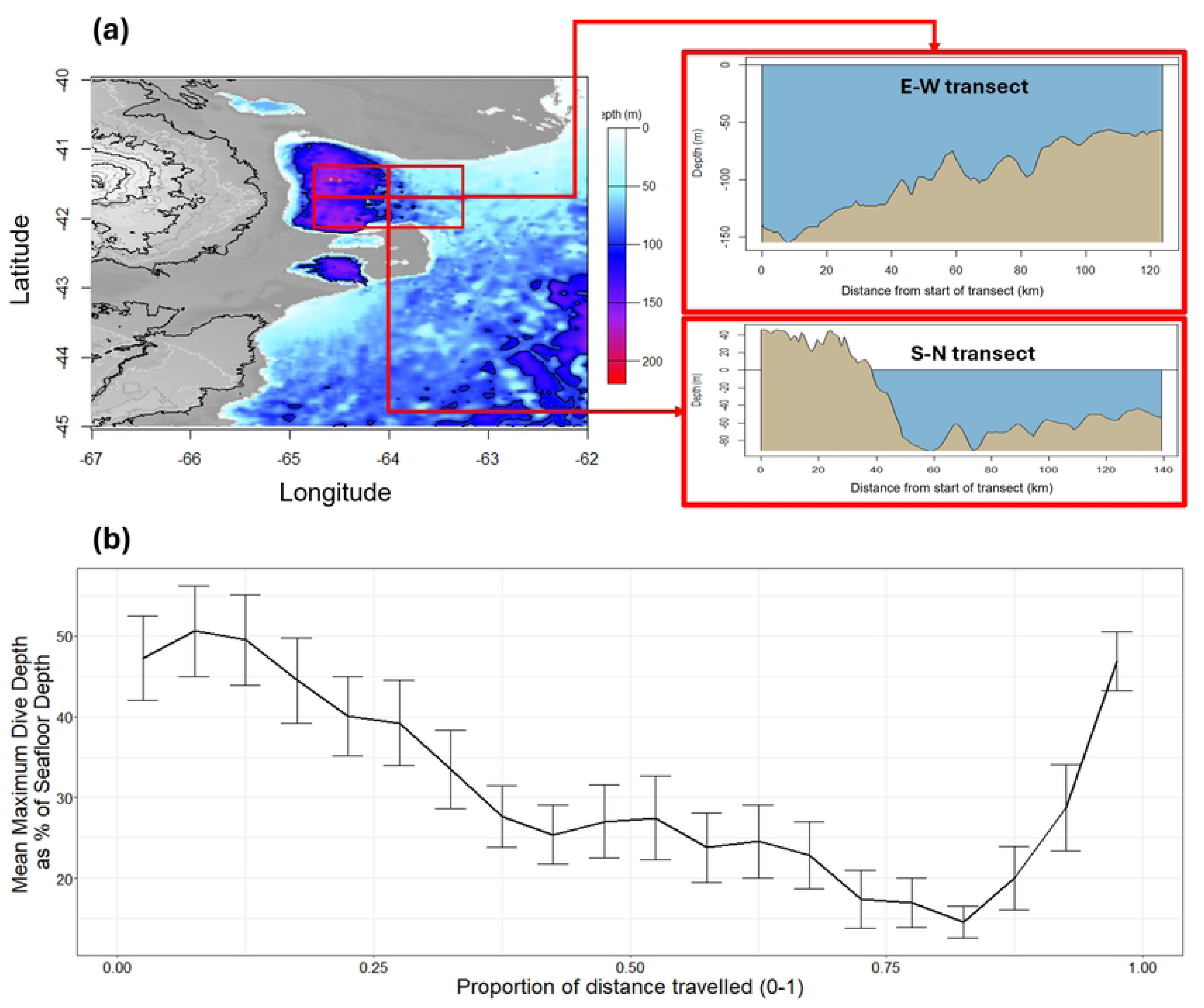
Bathymetry estimates of the San Matías Gulf. (a) Estimated seafloor depths in the region encompassing all penguin foraging journeys. (b) Mean maximum dive depth as a percentage of seafloor depth (± 1 SE) across proportions of total distance travelled during return trajectories. Percentage values were averaged per 0.05 increments of total distance travelled.

**Figure S7.**
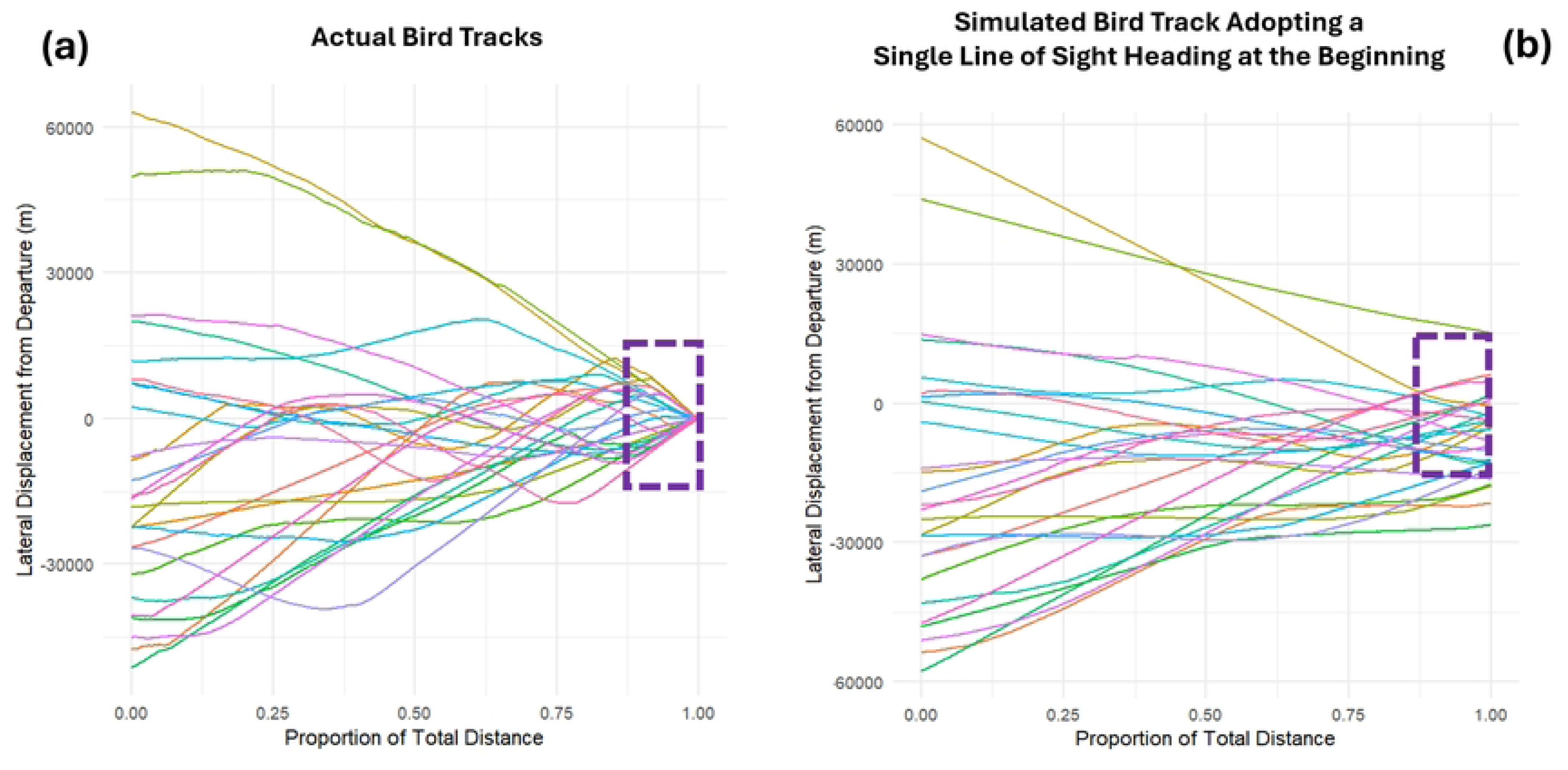
Lateral Displacement of Penguins Relative to Departure Point During Return Journeys. This figure illustrates the lateral displacement of penguins from their departure point at the colony during their return journeys. The y-axis represents lateral displacement (positive values indicate eastward movement; negative values indicate westward movement), and the x-axis represents the proportion of the total distance travelled (from 0 to 1). *(a)* Actual penguin data: Each line depicts an individual penguin’s lateral displacement over time. *(b)* Simulated scenario: Each penguin follows a fixed line-of-sight heading toward the colony without adjusting for current-induced drift effects. For both real and simulated penguins, starting coordinates were used as baselines to calculate absolute lateral displacement. The purple dashed boxes highlight displacement within 15 km of the colony during the final stages of the of the total distance travelled.

**Figure S8.**
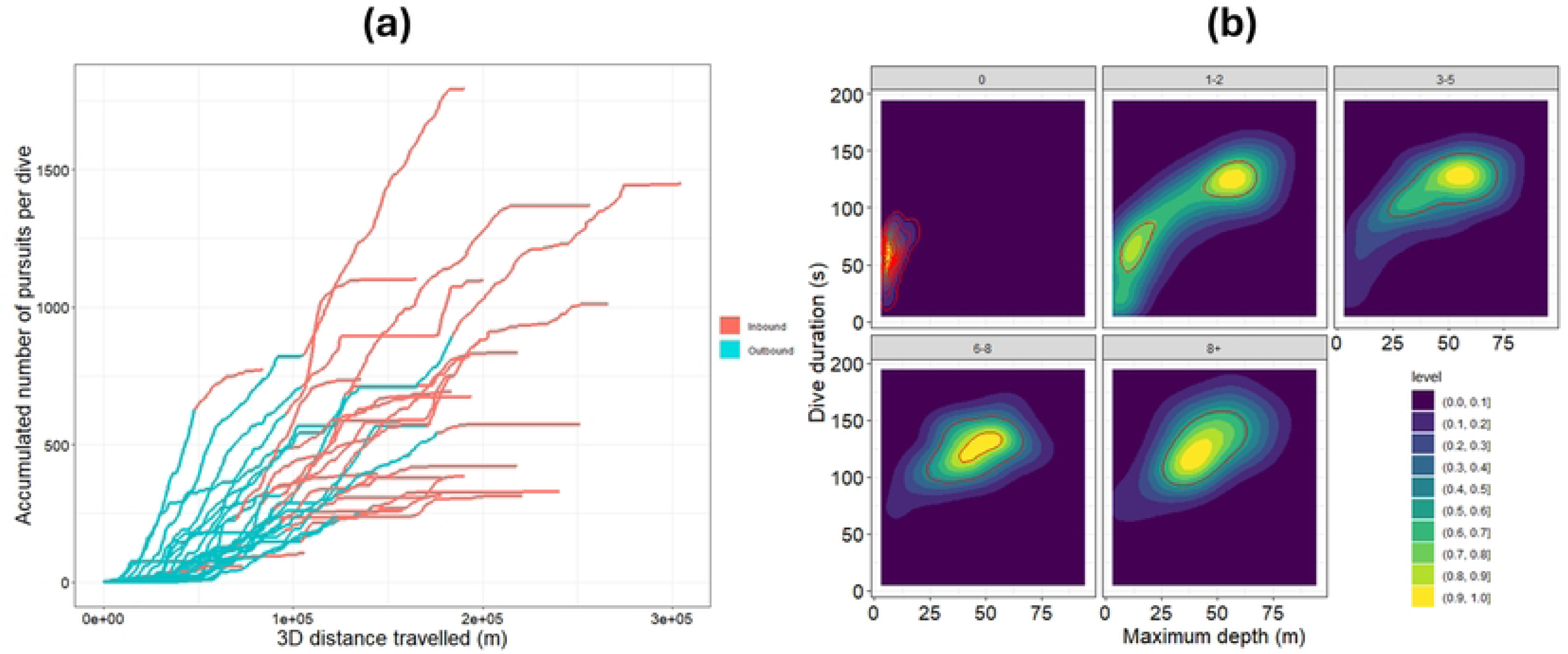
Prey Pursuits and Dive Depth vs Dive Depth. (a) The cumulative number of prey pursuits as a function of 3D distance travelled across each bird’s dead-reckoned path. Undulations in the dive profile were used as a proxy for prey pursuits, following methods outlined in Simeone and Wilson [14]. (b) The plots are faceted based on the number of prey pursuits per dive, illustrating how dive behaviour varies with foraging activity.

**Figure S9.**
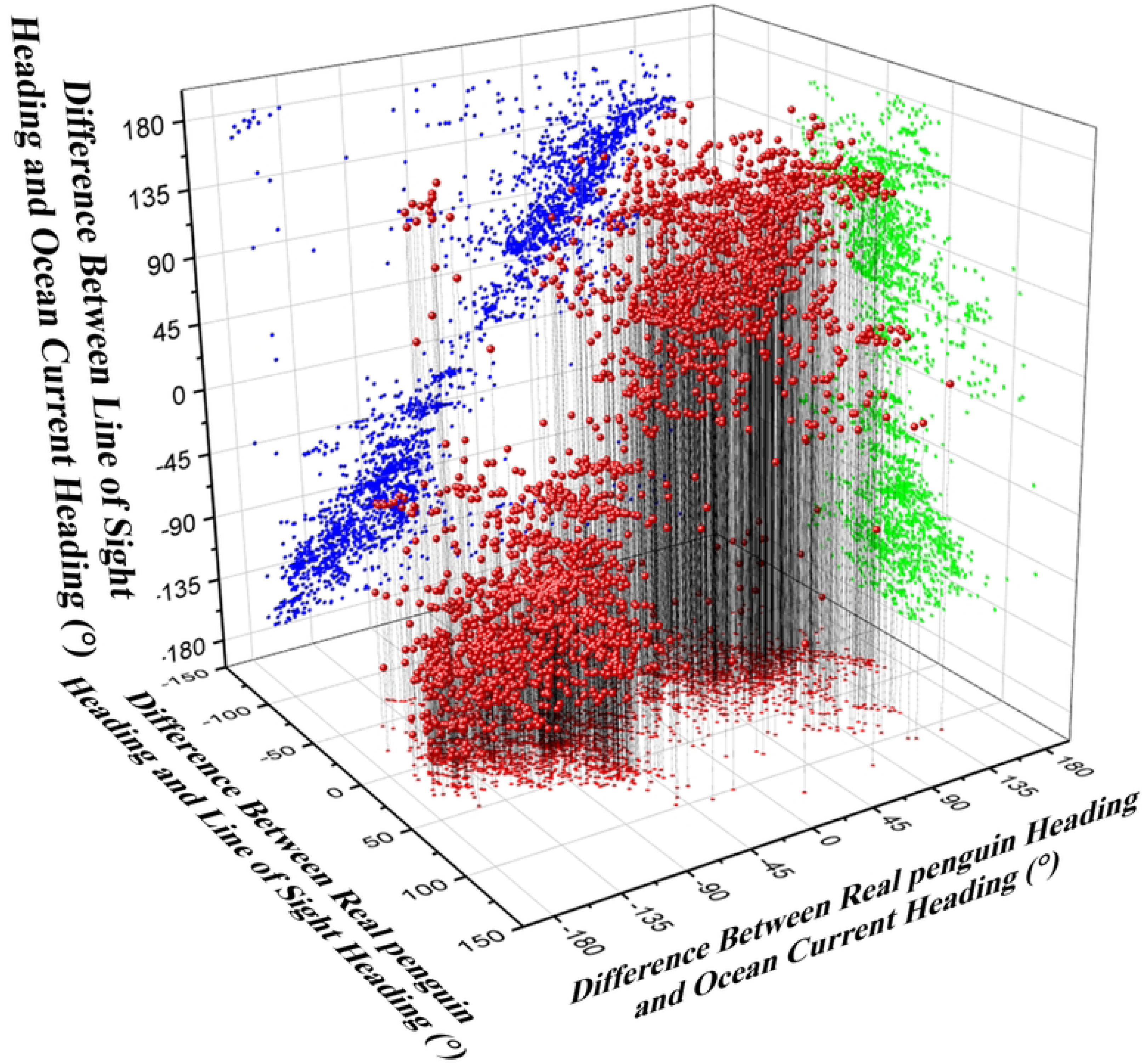
3D Scatter Plot of Angular Heading Differences Among Real Penguin, Line-of-Sight, and Ocean Current Directions. This figure illustrates the angular differences between the real penguin’s travel vector heading (relative to the water), the line-of-sight heading to the colony, and the ocean current heading. **X-axis (red x- y projection):** Angular difference between penguin heading and line-of-sight heading to the colony. **Y-axis (green z-x projection):** Angular difference between penguin heading and ocean current heading. **Z-axis (blue y-z projection):** Angular difference between line-of-sight heading to the colony and ocean current heading. Each red data point represents the mean angular difference calculated per 0.01 bin of each penguin’s proportion of distance travelled.

**Figure S10.**
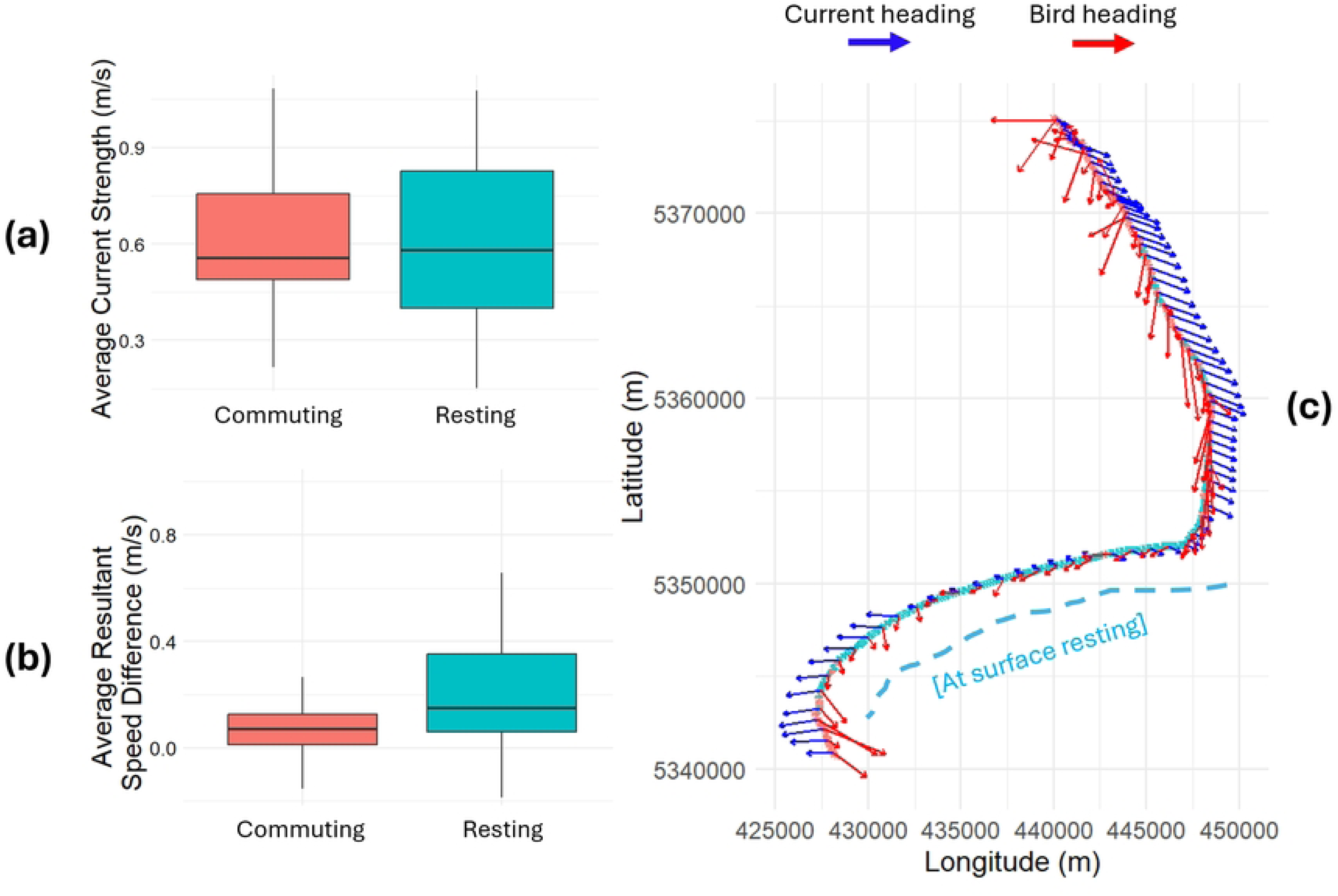
Comparison of Ocean Current Strength Conditions During Commuting and Resting Phases of Penguins’ Return Journeys. Resting was defined as surface periods ≥ 60 s and commuting as all times when birds were at depth (diving). (a) shows the average current speed (m/s) during commuting and resting phases, while (b) depicts the differences in resultant speeds (post-current integration vs. pre-current integration) for the two activity types. Boxplot (boxes encompass the 25∼75 % interquartile range, horizontal bars reflect the median and whiskers extend to 1.5 * Interquartile range) were constructed using the mean values per bird across activity types. (c) Presents an example of a penguin’s return journey exhibiting extended flow-assisted surface resting during slack current conditions. Red arrows denote the Actual bird headings and blue arrows denote the current direction. Arrow lengths are proportional to the speed of travel (both for the bird and the current).

**Figure S11.**
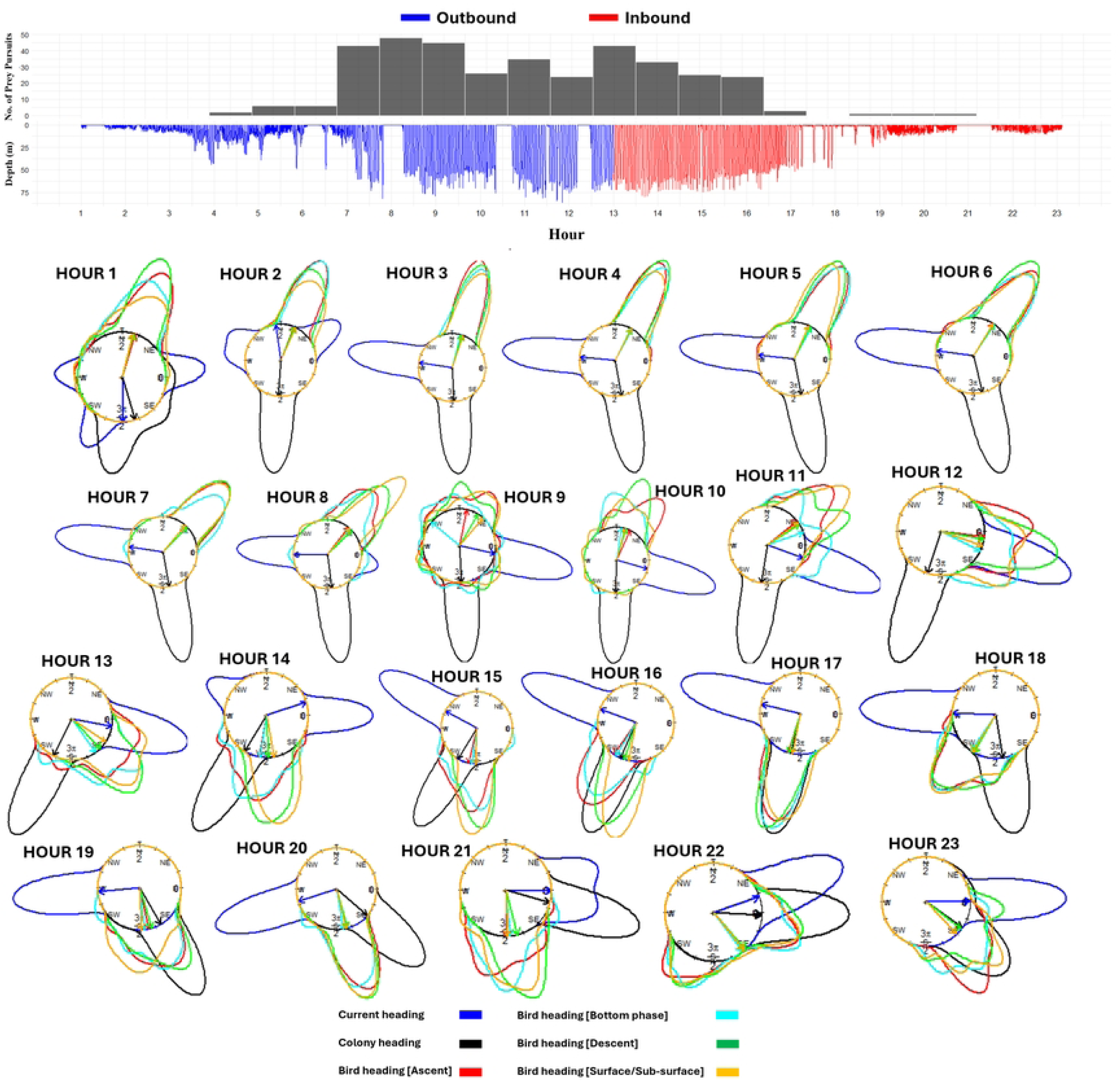
Example Data Showing Changes in Depth, Prey Pursuits, and Heading Distributions of Mean Circular Headings per Dive and Dive Phase for a Single Penguin. The **Top Panel** shows the number of prey pursuits (black bars) and depth profiles over time, with outbound dives indicated in blue and inbound dives in red. The **Bottom Panel** presents circular rose plots showing the mean headings per dive phase, color-coded as follows: descent (green), bottom phase (cyan), ascent (red), and surface/sub-surface (orange). It also includes the mean LoS heading towards the colony (black) and the ocean current heading (blue). Grand hourly means of these values are represented by arrows inside the circular plots.

## Notes

### Competing Interest Statement

The authors have declared no competing interest.

